# High-throughput targeted amplicon screening tool for characterizing intrahost diversity in *Staphylococcus aureus* directly from sample

**DOI:** 10.1101/2024.09.27.615532

**Authors:** Tara N. Furstenau, Ryann Whealy, Skylar Timm, Alexander Roberts, Sara Maltinsky, Sydney Wells, Kylie Drake, Ann Ross, Candice Bolduc, Talima Pearson, Viacheslav Y. Fofanov

## Abstract

A significant proportion of people are asymptomatic carriers of *Staphylococcus aureus* (SA), an important risk factor for development of opportunistic infections. SA colonization is dynamic, appearing and disappearing, with strains evolving and potentially shifting in composition over time and between body sites. These changes make detection challenging and the numerous potential sources of reintroduction from other people and even other body site reservoirs preclude efficient efforts to prevent transmission and spread. Identifying typical sources is therefore critical for mitigation. Whole-genome sequencing (WGS), ideally of multiple colonies from multiple body sites, is the gold standard for characterizing SA strains and confirming transmission. However, this is often too resource-intensive for initial assessments of transmission and not feasible for large-scale studies involving various body sites from multiple individuals over time. To address these challenges, we developed a low-cost, custom, species-specific amplicon sequencing (AmpSeq) assay, optimized to provide high resolution discrimination of SA genotypes directly from samples.

We tested this approach on a subset of samples that were a part of a large-scale longitudinal study of SA carriage. Oral and nasal samples were collected from 9 participants every two weeks for up to 18 weeks and qPCR positive samples were analyzed using our AmpSeq assay directly from the sample without culturing. The longitudinal sampling strategy enabled us to characterize changes in SA colonization patterns over time, detect potential strain mixtures, and identify rare variants that may serve as signatures of transmission between different body sites or among individuals. Without using WGS, we were able to rapidly eliminate the possibility of transmission between sampled residents. Participants that had positive oral and nasal samples had no fixed SNP differences between the two body sites, suggesting likely within-person spread. In these cases, we were able to infer the most likely direction of spread (nasal to oral sites) by analyzing segregating rare variants. While WGS can be used to provide higher resolution to colonization patterns and validate these findings, our amplicon sequencing approach offers a rapid, cost-effective, direct-from-sample method for species-specific screening intended for population-level characterization that allows researchers to characterize strain types, identify or eliminate likely transmission cases, and identify potential reservoirs before resorting to more expensive WGS methods.

**Authors summary:** Colonizing opportunistic pathogens like *Staphylococcus aureus* present a unique challenge for disease study because rather than causing acute infections upon transmission, they persist asymptomatically for long periods of time allowing the bacterial population to evolve and differentiate. Characterizing the diversity within these populations is important for choosing correct treatments, quantifying the risk of horizontal gene transfer, and understanding paths of transmission between people and spread to different body sites. The gold-standard approach for characterizing population diversity is through culturing and whole-genome sequencing of multiple colonies per sample which is labor-intensive and expensive for any large-scale study. Using a custom-designed species-specific amplicon sequencing assay, we offer a cost-effective method for characterizing the diversity in *Staphylococcus aureus* populations directly from samples without the need for labor-intensive culturing or whole-genome sequencing. Our small-scale study highlights how this method provides a scalable tool for large epidemiological studies ideal for systematically exploring broader patterns of carriage and transmission.

## Introduction

*Staphylococcus aureus* (SA) is a significant public health concern due to its ability to cause a wide range of infections, from mild skin infections to life-threatening conditions such as bacteremia, pneumonia, and endocarditis [1–3]. The emergence of antibiotic-resistant strains, such as methicillin-resistant SA (MRSA), has further heightened the risk associated with SA, complicating treatment options and leading to higher morbidity and mortality rates [4–6].

SA lives in close association with humans and can persist asymptomatically for long periods of time. Various body sites are frequently colonized, particularly the skin and mucosal surfaces and opportunistic infections occur when the bacteria penetrate these barriers [7–10]. While infections can be due to extrinsic sources, there is a tight link between carriage and self-infection [2,7,10–15]. Much remains unknown about carriage patterns, but as detection methods improve, estimates of carriage frequencies have increased with a recent estimate suggesting that >65% of the population are carriers [6,16]. Carriage is most often identified in the anterior nares, but it is often found in the oropharynx, skin, and other mucosal surfaces [17]. SA colonized body sites can serve as reservoirs for spread to other body sites, endogenous infections, and transmission to others [11,18,19]. Longitudinal studies assessing changes in presence/absence, pathogen quantities, and genotypes can produce valuable insights into carriage, but have been severely limited due to a lack of high-throughput and high-resolution methodologies [18,20–27]

Understanding the dynamics of bacterial populations within hosts is crucial for understanding carriage and transmission patterns. Compared to the short timeframe of an acute infection, the persistence of SA during long-term carriage allows for the accumulation of mutations that can be lost or propagated through genetic drift or selection [28–30]. Additionally, co-colonization of multiple strains, either within or between different body sites, occurs relatively frequently [31–33] and can increase the potential for horizontal gene transfer and promote the spread of antibiotic resistance and virulence genes [34–36]. Failure to capture intrahost diversity can complicate the diagnosis, treatment, and tracking of infections, as different strains within the same host may respond differently to antibiotics or evade immune detection. Characterizing pathogen genetic diversity can also help in identifying key reservoirs and tracing transmission pathways [21,37] that may be missing or incomplete if only a small number representative isolates are compared. By comprehensively studying intrahost diversity, we can better understand the evolution of SA within hosts, improve infection control strategies, and develop more effective interventions to prevent the spread of this pathogen.

Whole genome sequencing (WGS) is currently the gold-standard method for characterizing SA strains because it provides comprehensive genomic information. However, accurately capturing the full complexity and heterogeneity of a colonizing SA population requires sequencing multiple isolates from each sample, as single isolates are not representative of the entire population’s diversity [32,38]. Additionally, sampling from multiple body sites is necessary to identify all potential reservoirs of SA within a host while sampling multiple timepoints is necessary for understanding how the colonizing population changes or evolves over time. WGS approaches provide invaluable insights, but also present significant challenges. The large number of samples needed to systematically characterize longitudinal within host diversity and necessity of culturing, isolating, and sequencing numerous colonies from each sample, makes WGS approaches both time-consuming and cost prohibitive.

PCR-based molecular typing methods like MLST [39] that are adapted to a next-generation sequencing platform offer a more economical alternative to characterizing population variation. PCR-based approaches can be applied directly to the sample without the need for culturing and can provide very high coverage of population variation within the targeted regions even with low concentrations of starting DNA. Unfortunately, the small number of housekeeping genes targeted by traditional MLST provide comparatively low discriminatory resolution that is not optimal for either confirming or refuting transmission events with any degree of confidence and the genes/primers are often not species specific making it difficult to use for population level characterization. However, if a larger number of strategically selected targets were queried, targeted amplicon sequencing (AmpSeq) could offer a cost-effective way to genotype the sample and characterize the diversity within bacterial populations.

We are currently conducting a large-scale longitudinal study of SA carriage and transmission involving the collection of oral and nasal swabs every two weeks. Our goals are to 1) characterize SA carriage rates, 2) identify circulating strain types, 3) track how colonizing strains are replaced or evolve over time, 4) characterize the frequency and dynamics of co-carriage of multiple strains (within or between body sites), 5) determine if there are associations between strain types and body sites 6) determine which body sites are more likely to act as reservoirs for the spread of SA, and 7) detect possible transmission between residents. Answering these questions using WGS on tens of thousands of isolates (assuming multiple isolates per positive sample) is costly and inefficient. Therefore, we need a rapid, high-throughput, cost-effective method to characterize the colonizing population and help screen for likely reservoirs and transmission pairs so that WGS resources can be reserved for validating and providing additional population genetic details for only these candidates. Here we describe a custom AmpSeq assay designed to address these challenges.

## Methods

### Sample collection

Samples were collected every two weeks from residents of three long-term care facilities in the Phoenix Metropolitan area. Informed written consent was received from each participant and the project was approved by the Northern Arizona University Institutional Review Board (Project 1766728-3). Swabs of the anterior nares and mouth were self-administered under the supervision of study staff to ensure consistency. This method has been highly successful in detecting SA in previous studies [16,40,41]. After sampling, swabs were stored in 1mL of Liquid Amies medium and stored in a -20°C freezer before processing.

### DNA extraction and *S. aureus* detection

DNA was extracted from the collection media using an Applied Biosystems MagMax DNA Multi-Sample Ultra 2.0 extraction kit on a Thermo Fisher KingFisher Flex instrument. The SaQuant qPCR assay [16,42] was run in 10 µl reactions using 5ul of Applied Biosystems TaqMan Universal PCR MasterMix, 1 µM of forward primer, 1 µM of reverse primer, 200 nM of TaqMan TAMRA probe, and 1 µl of template. Thermo cycling conditions were as follows: hot start TaqMan activation (10 minutes at 95°C), followed by 40 cycles of denaturation (15 seconds at 95°C) and extension (1 minute at 57°C).

### Target and primer design for multiplexed amplicon sequencing assay

Our custom amplicon sequencing assay was designed to target genomic sites that were conserved across strains but contained polymorphisms that maximized strain differentiation within SA. The sites were bioinformatically confirmed to be species specific so that similar loci in other common commensal *Staphylococcus* species would not amplify and skew variant detection. A set of 961 SA reference genomes were downloaded from the RefSeq database [43] using the ncbi-genome-download tool (https://github.com/kblin/ncbi-genome-download). The genomes were aligned to the NCTC 8325 reference genome (NC_007795.1) using NUCmer v3.1 [44] and SNPs were identified using NASP v1.2.1 [45] with defaut parameters after masking duplicated loci and regions that had high similarity with reference genomes from 12 other *Staphylococcus* species commonly found in human samples: *S. epidermidis, S. haemolyticus, S. saprophyticus, S. hominis, S. warneri, S. capitis, S. simulans, S. cohnii, S. xylosus, S. saccharolyticus*, and *S. lugdunensis* [46–48]. VaST [49] was used to identify groupings of SNPs within 100bp windows that maximized strain-level resolution among the reference genomes. Primers were designed to amplify the 27 selected targets (Supplemental Table 1) and were optimized to work together in a single multiplex PCR (i.e., minimizing the number of primer interactions and ensuring similar melting temperatures). The primers were synthesized with forward (UT1: 5’-ACCCAACTGAATGGAGC -3’) and reverse (UT2: 5’-ACGCACTTGACTTGTCTTC -3’) universal tails to facilitate index ligation prior to pooling for multiplexed sequencing.

### Amplicon Sequencing

The KAPA 2G Fast Multiplex PCR Master Mix was used with 27 primer pairs at 0.2 µM concentration, with 5ng of template DNA and enough water to bring the reaction mixture to 25µL. The PCR thermocycling conditions consisted of an initial denaturation at 95°C for 3 minutes, followed by 35 cycles of denaturation at 95°C for 15 seconds, annealing at 60°C for 30 seconds, and extension at 72°C for 1 minute and 30 seconds and a final extension at 72°C for 1 minute. The PCR products were then prepared for sequencing using our standard amplicons sequencing protocol [50] and were sequenced on the Illumina MiSeq platform to generate 150 bp paired end reads.

### SNP calling and rare variants

The primer sequences were removed from both the forward and reverse sequencing reads using Cutadapt v4.9 [51]. The reads were then processed using Trimmomatic v2.9.1 [52] with the following parameters for quality control: LEADING=3, TRAILING=3, SLIDINGWINDOW=4:15, and MINLEN=36. Reads were aligned to the reference sequence (NC_007795.1) using BWA-MEM v2.4.0 [53,54]. SNPs and rare variants were called using LoFreq v2.1.5 [55] after running the ‘viterbi’ command to correct alignment errors. Consensus SNPs were called if there were at least 10 reads covering the site and the majority (>70%) of reads supported the call.

A group of high confidence rare variants was generated by collecting loci where LoFreq called a low frequency variant (<2%) in at least one sample. For samples where a rare variant was not called at a position with LoFreq, the missing data was filled with pileup allele frequencies generated using BCFtools v1.15.1 [56] with a base quality cutoff of 20, a mapping quality cutoff of 20, and a minimum coverage of 10 base pairs. Only genome ranges within each amplicon, excluding the primer regions, were included in analyses. Because some rare variants were likely a result of errors introduced through PCR and sequencing, we focused on counting rare variants that were detected in samples at multiple timepoints to increase our confidence that true variants were detected. Rare variant allele frequencies were used to calculate Nei’s genetic distance [57] between the populations at different time points.

### Phylogenetic analysis

To identify the sequence types of the samples, we drew a minimum spanning tree using Consensus SNPs from our target sites in both the samples and reference genomes using Grapetree v2.1 [58] using the MSTreeV2 method. The sequence types for the reference genomes were estimated using mlst v2.23.0 [39,59,60] and our samples were assigned to a sequence type based on the sequence type of the reference that they clustered with. PAUP v4.0a [61] was then used to draw a maximum parsimony tree using only our samples and the consensus SNPs identified within them.

## Results

### Sequenced Samples

The samples that were sequenced using the AmpSeq assay were from 9 different participants from three long-term care facilities. These participants were selected because they had multiple consecutive SA positive samples (via qPCR), and four participants were positive in both the oral and nasal samples, providing insights into longitudinal carriage and interactions between body sites. The positive samples selected for AmpSeq are indicated in Figure 1. For three participants (ff5b0b, 45e796, and 85b498), the sequenced samples spanned 18 weeks (Figure 1). Overall, 69 samples were sequenced, 55 were nasal and 14 were oral samples. The average depth of coverage per position was 716 (sd=492) across all samples with a minimum of 65 and a maximum of 2,847. The coverage for the nasal samples was generally higher (mean=827, sd=480) than the oral samples (mean=279, sd=232). Of the 27 amplicon targets, 22 had consistent coverage with at least 50/69 samples having at least 50x coverage and a minimum average coverage of 250x per locus (Supplemental Figure 1). The sample metadata, average coverage, and SaQuant results are available in Supplemental Table 2.

**Figure 1.**
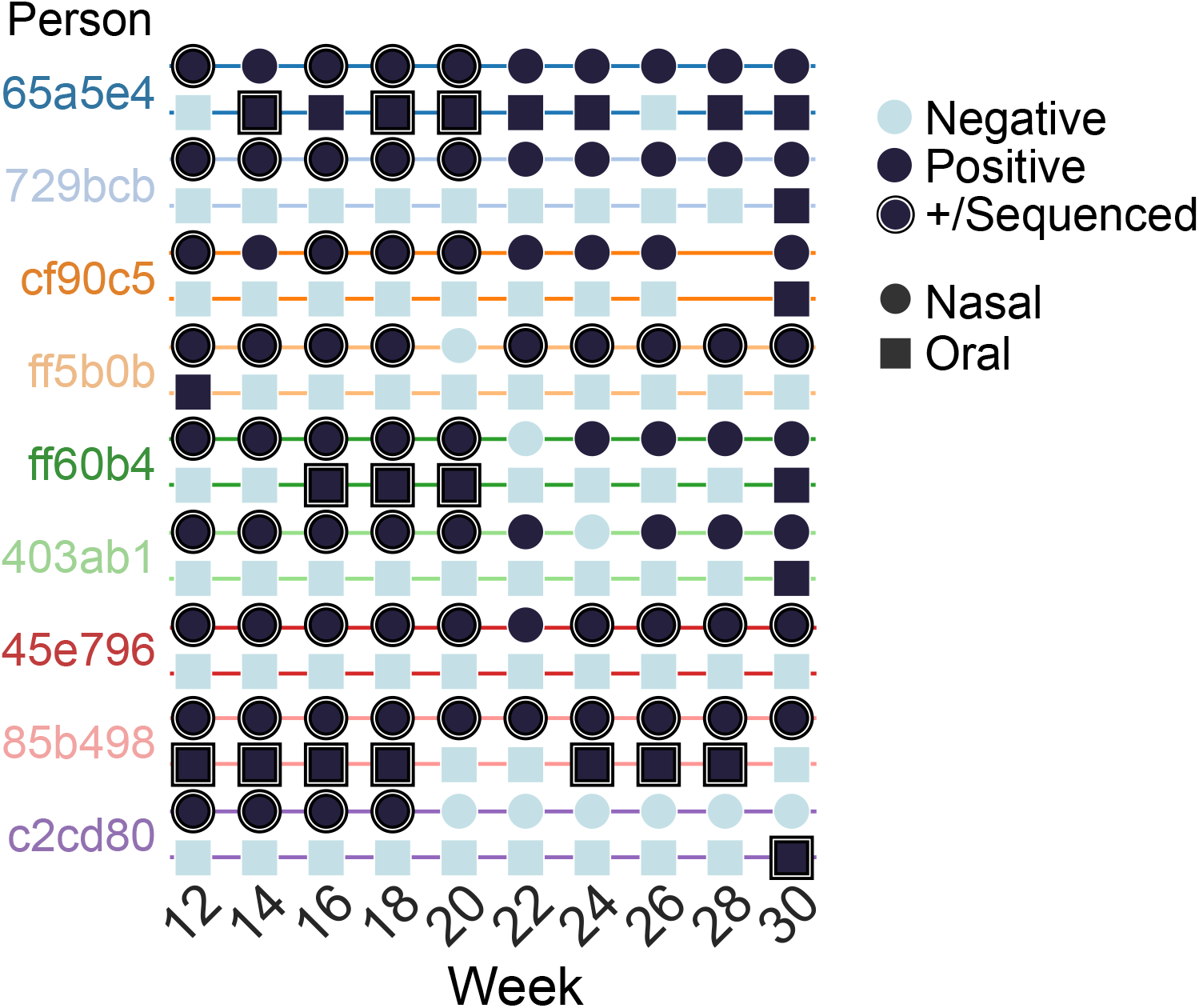
Timeline of *S. aureus* carriage among nine long-term care facility residents. Nasal (circles) and oral (squares) samples were collected from participants every two weeks and tested for SA using qPCR. Positive samples are indicated in dark blue, positive and amplicon sequenced samples are indicated in black.

**Supplemental Figure 1.**
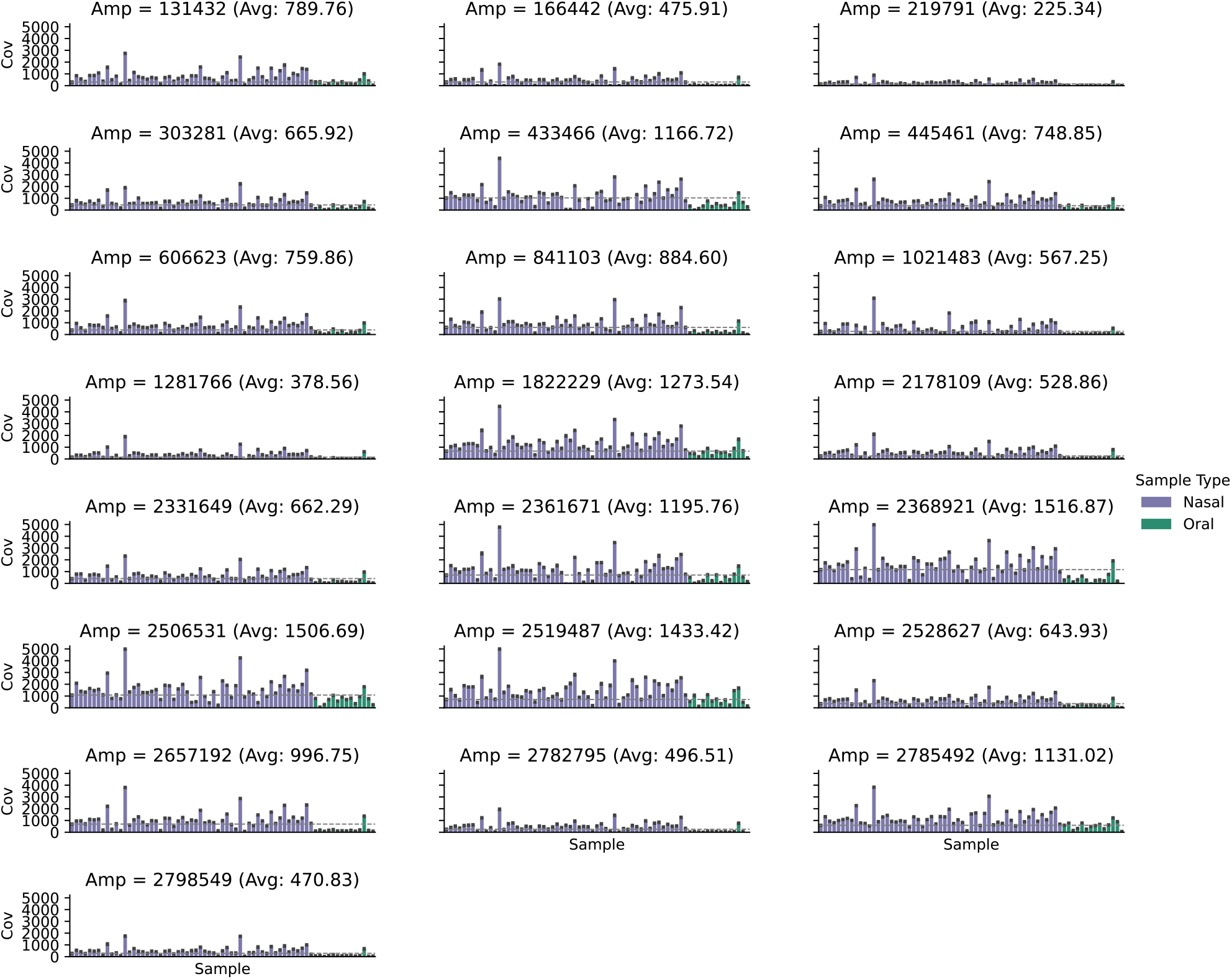
Consistent coverage was achieved at 22 of the amplicon targets. The figure shows the coverage for each sample for each amplicon target. The average depth of coverage across all samples was 716 (sd=492). Each amplicon had at least 50/69 samples with greater than 50x coverage. The minimum average coverage for an amplicon was 255x for amplicon 219791. The coverage for the nasal samples (purple) was generally higher (mean=827, sd=480) than the oral samples (green, mean=279, sd=232).

### Strain typing

Our selected targets provide high resolution discrimination among sequence types and the strains found within the long-term care facilities were highly diverse. Across all samples, a total of 1,806 sites were identified that had a majority SNP allele that was different than the reference sequence and 133 of the sites were parsimony informative. Figure 2 shows the placement of our samples among reference sequences with known sequence types. The strains colonizing different individuals are diverse, with 9 genotypes representing 7 different sequence types.

**Figure 2.**
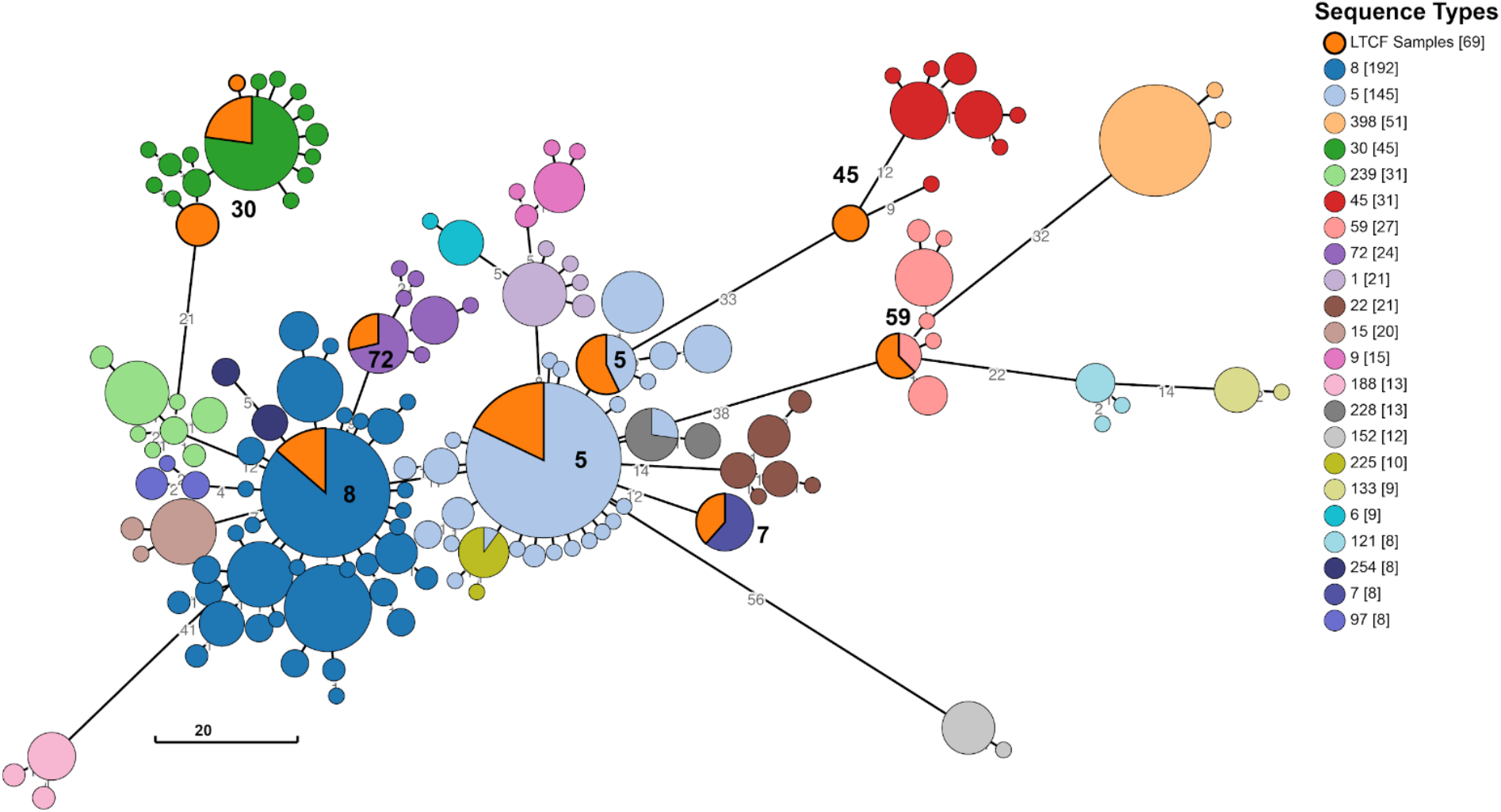
The selected targets offer clear differentiation between sequence types and high-resolution discrimination within sequence types. Minimum spanning tree comparing the relationships between reference sequences with known sequence types and the samples from our study. The minimum spanning tree was drawn using the consensus SNPs identified within the amplicons of our AmpSeq assay. The predicted sequence types of our samples (shown in orange with a bold outline) is indicated next to the nodes.

Figure 2 also shows how our selected targets provide resolution within sequence types, particularly identifying multiple genotypes within the common ST5 and ST8 groups. Within each individual, the samples collected at different timepoints and at both body sites were identical when looking at the majority (consensus) alleles at each site (Figure 3). Analysis of the amplicon sequences showed no indication of mixed haplotypes within a sample, so no cases of co-colonization were detected. Across individuals, there were at least 2 SNPs differentiating the most closely related genotypes out of a total 3,476 possible positions covered by the amplicons. Two people carried strains that were predicted to be ST30 and two carried strains that were likely ST5, however in each of these cases, the resulting AmpSeq genotype was different between people. In both cases the participants were from different facilities so there was a low likelihood of interactions that may have resulted in transmission. All other individuals carried unique AmpSeq genotypes that formed distantly related monophyletic groups.

**Figure 3.**
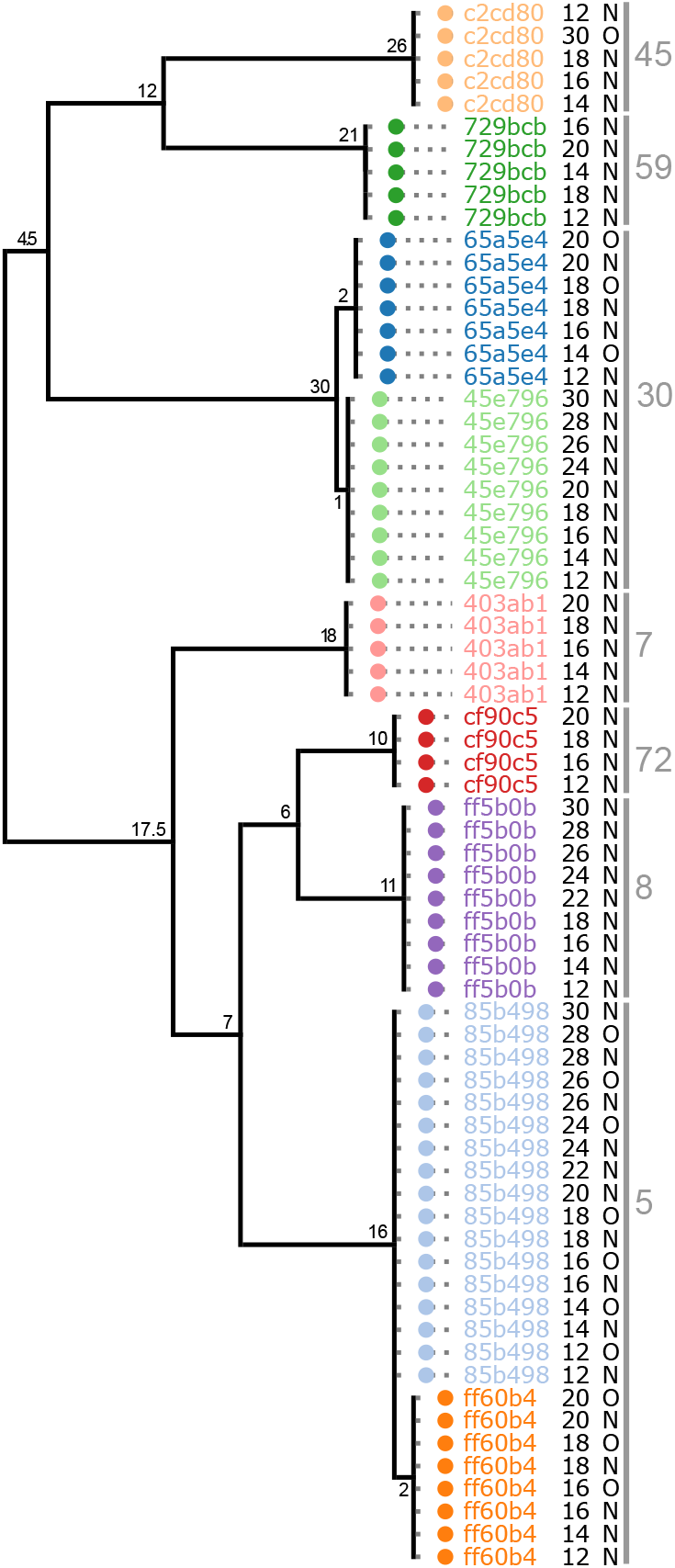
Carriers maintained the same strain over time and shared the same strain at both body sites; there was no indication of transmission between individuals. The parsimony tree indicates the number of SNP differences between the samples on the branches. Samples from the same individual are color coded and labeled in the first column, the week that the sample was collected is labeled in the second column, the body site from which the sample was collected is labeled in the third column, and the final column indicates the predicted sequence type for each clade.

### Low frequency variants maintained in populations over time

With deep sequencing of the amplicons, we identified low frequency variants (<2%) at 1,804 genomic positions. These represent point mutations that are segregating in the population that have not yet been fixed or removed. While some variants were lost over time or undetected, many were maintained in each population throughout the duration of sample collection (Figure 4). The time between samples was not a significant predictor of genetic distance (linear regression, p = 0.206) while average coverage was significant (p < 0.001). This suggests that rare variants were more likely undetected rather than lost from the population, further supporting the longevity of variants in the populations. The average frequency of rare variants detected in the nasal samples was 0.0029 (sd=0.0016) while the average frequency for the oral samples was 0.0041 (sd=0.0024). This difference is expected because the higher coverage in the nasal samples allowed us to detect variants that were present at lower frequencies.

**Figure 4.**
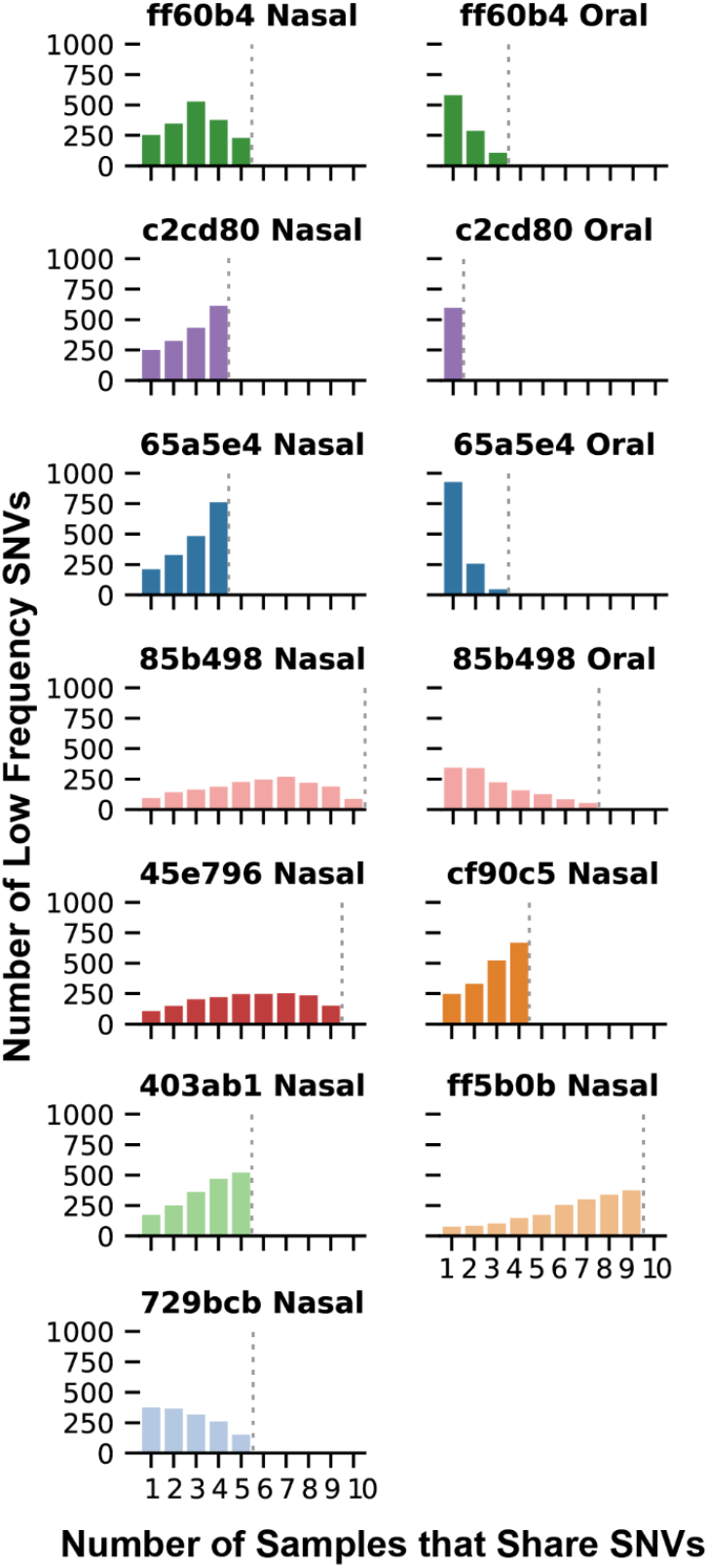
Many rare single nucleotide variants persisted in populations across all time points. The figure shows the number of low frequency variants (<2%) that are shared by a given number of samples from the same individual (x-axis). The dashed grey line indicates the total number of samples analyzed per individual and body site.

### Comparing diversity between nasal and oral samples

On average, the nasal samples had greater depth of coverage and lower Ct values (mean=23, sd=5.2) compared to the oral samples (mean Ct=30, sd=10.2). The consistently lower DNA quantities in the oral samples suggests that the population size may be smaller in the mouth. Additionally, the accumulation of low frequency variants with increasing depth of coverage plateaus more rapidly in the oral populations than the nasal populations indicating that most of the sample diversity was captured with the achieved depth (Figure 5). Figure 6 shows that only a few low frequency variants were shared among multiple oral samples that were not also shared with the nasal samples. The reverse, however, is not true for the nasal samples which share many low frequency variants that were not shared in the oral samples. Combined, this evidence suggests that the nasal populations are more established and are likely to be the source of the oral populations.

**Figure 5.**
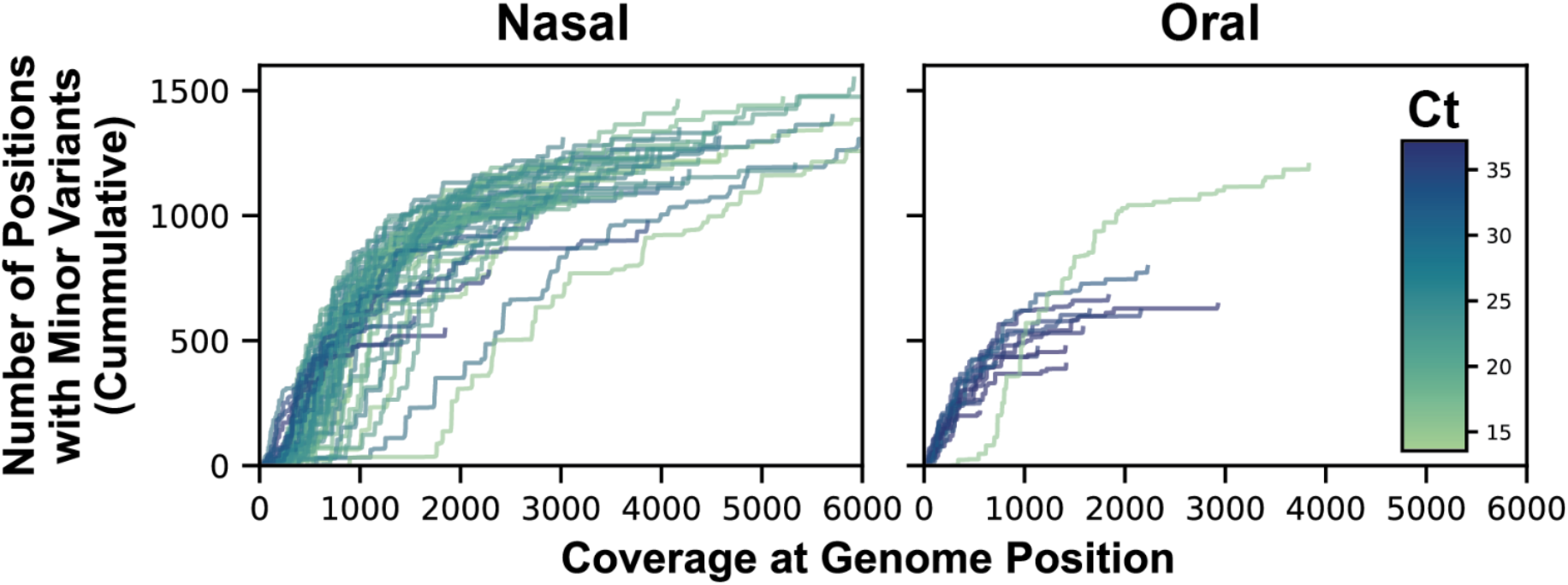
DNA quantity and accumulation of low frequency variants with coverage depth suggests that the effective population size in nasal samples is larger than oral samples. The oral samples generally showed higher Ct values than the nasal samples and the accumulation of low frequency variants with coverage plateaus more rapidly suggesting that most of the diversity was captured.

**Figure 6.**
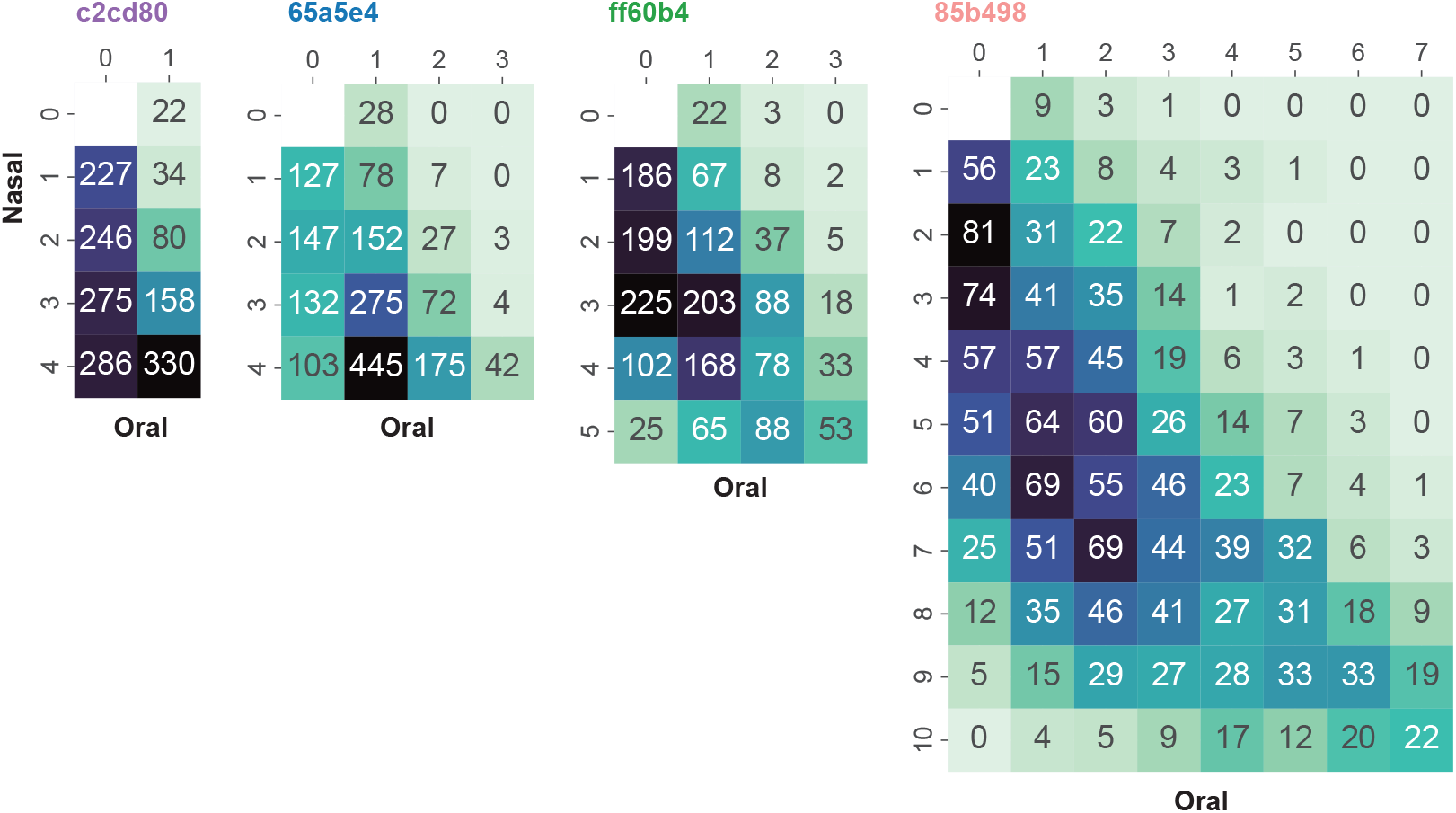
The nasal populations are the likely source of spread to the oral populations. The heatmap shows the number of low frequency variants that are shared between a given number of oral samples (x-axis) and a given number of nasal samples (y-axis). Few low frequency variants were shared among multiple oral samples that were not also shared with the nasal samples. The reverse is not true for the nasal samples which share many low frequency variants that were not detected in the oral samples.

For individuals 65a5e4 and ff60b4, the nasal populations were established prior to the oral populations by at least two and four weeks, respectively (Figure 1). Based on the number of rare variants the oral samples share with the nasal samples, it seems likely that the nasal population was the source for colonization of the mouth, with frequent introductions. For individual 85b498, four oral samples were positive followed by two negative samples in weeks 20 and 22 and then three more positives. The source of the re-established oral population could have been either the nasal reservoir which persisted through weeks 20 and 22, or a severely reduced oral population that persisted below our limit of detection. Our results suggest that the former is more likely because, if a bottleneck occurred as suggested by the later, we would not expect as many variants to be shared with as many of the nasal samples and we would expect more variants shared among the oral samples. Individual c2cd80 had positive nasal samples from week 12 to week 18, then oral and nasal samples were negative until a positive oral sample was detected at week 30. The one oral sample had only 22 (3.5%) low frequency variants that were not identified previously in the nasal samples and 330 (53%) that were shared with all four of the earlier nasal samples. Again, this may suggest that the source population for the mouth was either a nasal population that persisted below the level of detection or perhaps colonization was established through an unsampled body site or retransmission from an individual (unsampled) that was also the source of the initial nasal colonization. Nonetheless, the large number of variants shared between all the nasal and the oral sample, is suggestive of a nasal source and the timing of the positive oral samples suggest that the oral populations likely receive frequent introductions from nasal populations.

## Discussion

### Characterizing SA carriage

This study was an initial test on a small subset of samples to establish the utility of our AmpSeq approach in characterizing SA carriage for large-scale studies. In our small subset of 9 participants, we showed that they carried 7 distinct sequence types (9 distinct AmpSeq genotypes) and that the colonizing strains were stable for up to 18 weeks. Participants that were colonized in both the nose and the mouth tended to carry the same strain at both body sites and there were no cases of co-carriage of multiple strains. Although these results cannot be extrapolated for the whole long-term care facility population, it does demonstrate that this AmpSeq approach offers a cost-effective, yet high-resolution method for strain typing SA populations directly from samples without culture. When expanded to the analysis of our full large-scale study, this assay will allow us to understand population-level trends in carriage using high-powered analyses. It will enable us to characterize strain types circulating within the long-term care facilities, track how often strains change due to recolonization, estimate the rate of co-carriage, and establish associations between strain types and demographic data.

### Detection of putative transmission between individuals

Low-frequency variant diversity and sharing patterns can aid in identifying transmission sources and recipients as well as population dynamics over time. However, our AmpSeq assay does not provide sufficient resolution for definitively characterizing population dynamics and directionality of spread/transmission. For such a definitive assessment, we have developed a novel WGS approach that also leverages population-level data [38] for diversity and transmission analysis. Our AmpSeq method can be used to quickly eliminate potential transmission pairs that do not have matching genotypes. With this filter, expensive WGS verification of transmission can be reserved for only the most likely or high-priority cases. In the samples tested here, we show that our amplicon targets provide greater resolution than traditional MLST approaches, allowing us to differentiate between strains within a sequence type. As a result, although multiple pairs of individuals carried similar sequence types, we could determine that they were unlikely to be related due to multiple SNP differences detected among our amplicon targets.

### Detection of putative host reservoirs and sources of spread to other body sites

The nares are commonly reported to be the most frequent site of colonization and are a likely reservoir for transfer to other body sites [2,11,16,18,19,62–64], and our results are consistent with this body of literature. In individuals with positive samples at both body sites, the nasal samples were typically positive more frequently than the oral samples. We observed that nasal samples consistently exhibited higher DNA quantities [16], and they carried more diversity.

Additionally, the fact that many low-frequency variants found in nasal samples were also detected in oral samples, but not vice versa, indicates that the nasal population may be seeding other sites within the same host. The persistence of these low-frequency variants across multiple weeks further underscores the stability and central role of the nasal population in long-term carriage. These findings align with previous studies that have identified the anterior nares as a major reservoir for SA, playing a crucial role in both endogenous infections and the transmission to other sites within the body [62,63]. However, studies of genetically verified spread between sites tend to be smaller in scale and limited by either the cost of WGS or the low discriminatory power of MLST. Large-scale studies are lacking but are required to provide the statistical power necessary to fully support these contentions and establish consistent and actionable patterns of spread between body sites.

### Caveats

One of the goals of this study was to understand the microvariation that exists within colonizing SA populations and utilize this information to gain insights into population dynamics, evolution, and epidemiology. While PCR can amplify low-abundance sequences, it is also prone to introducing errors during amplification, which can create false-positive variants. This issue is particularly relevant when dealing with rare variants that occur at low frequencies, as distinguishing true biological variants from PCR- or sequencing-induced artifacts becomes difficult. Our study did not include technical replicates, which are typically used to identify spurious variants introduced during PCR amplification. Instead, we relied on the availability of multiple samples collected from the same individual across different time points to assess the persistence and reliability of rare variant detection. We found that many rare variants were consistently maintained across multiple samples for up to 18 weeks. The maintenance of low frequency alleles (~0.3% on average for the nasal samples) for long periods of time suggests that selection on these loci may be weak and/or the effective size of the colonizing population is high enough that these variants are not rapidly fixed or removed through genetic drift. True technical replicates would be required to determine accurate time frames for shorter lived true biological variants. But the fact that a non-trivial number of rare variants were maintained across all samples suggests that longer sampling intervals could be sufficient for tracking the dynamics of colonizing populations over time.

### Future Applications

The approach described in this paper has several potential applications for large-scale research studies and we intend to continue employing this strategy as we systematically explore broader patterns of carriage and transmission. Our approach is ideal for 1) cost effectively characterizing strain types carried by a large group of hosts and tracking changes in strain types over time 2) identifying multi-strain infections – targeted amplicon sequencing provides deep coverage of multiple targets which can reveal mixtures of haplotypes within a population 3) studies that require high statistical power, like associating strain types with certain host characteristics or body sites, 4) Identifying reservoirs for spread and transmission – shared rare variants can help establish sources of spread between body sites and transmission to other individuals and help quantify how much diversity is transferred between body sites and between hosts. Because we can amplify directly from the collection media without culturing, this approach produces rapid results. While this method is not intended to produce definitive results regarding genotype similarity or validate transmission cases, it serves as an efficient and inexpensive screening tool to identify samples of interest for further validation. This can significantly reduce the need for whole-genome sequencing to only a subset of samples thereby conserving resources while still obtaining high-resolution data.

## Conclusion

Our small-scale study demonstrates the utility of targeted amplicon sequencing for characterizing colonizing SA populations and detecting and tracking rare variants within SA populations. The findings provide valuable insights into the persistence of these strains and rare variants over time and offer practical guidance for future studies aimed at understanding bacterial diversity, sources of spread, transmission, and infection control. As we continue to refine this approach, it holds promise for broad applications across pathogen species in both clinical and epidemiological research, ultimately contributing to more effective strategies for controlling the spread of bacterial pathogens.

## Author statement

**Tara N. Furstenau:** *conceptualization, data curation, funding acquisition, investigation, methodology, visualization, writing – original draft, review, and editing*.

**Ryann Whealy**: *data curation, investigation, methodology, writing – original draft, review, and editing*.

**Skylar Timm**: *data curation, investigation, writing – review and editing*.

**Alexander Roberts:** *data curation, investigation, writing – review and editing*.

**Sara Maltinsky:** *data curation, investigation, writing – review and editing*.

**Sydney Wells:** *data curation, investigation, writing – review and editing*.

**Kylie Drake:** *data curation, investigation, writing – review and editing*.

**Ann Ross:** *investigation, resources, supervision, writing – review and editing*.

**Candice Bolduc:** *investigation, resources, writing – review and editing*.

**Talima Pearson:** *conceptualization, funding acquisition, supervision, methodology, writing – original draft, review, and editing*.

**Viacheslav Y. Fofanov:** *conceptualization, funding acquisition, supervision, methodology, writing – original draft, review, and editing*.

## Repositories

The amplicon sequencing reads have be deposited in NCBI’s Sequencing Read Archive under BioProject PRJNA1166327.

## Ethical approval

Informed written consent was received from each participant and the project was approved by the Northern Arizona University Institutional Review Board (Project 1766728-3).

