## Supplementary figures and images for "High-throughput targeted amplicon screening tool for characterizing intrahost diversity in *Staphylococcus aureus* directly from sample"

### Supplemental Figure 1

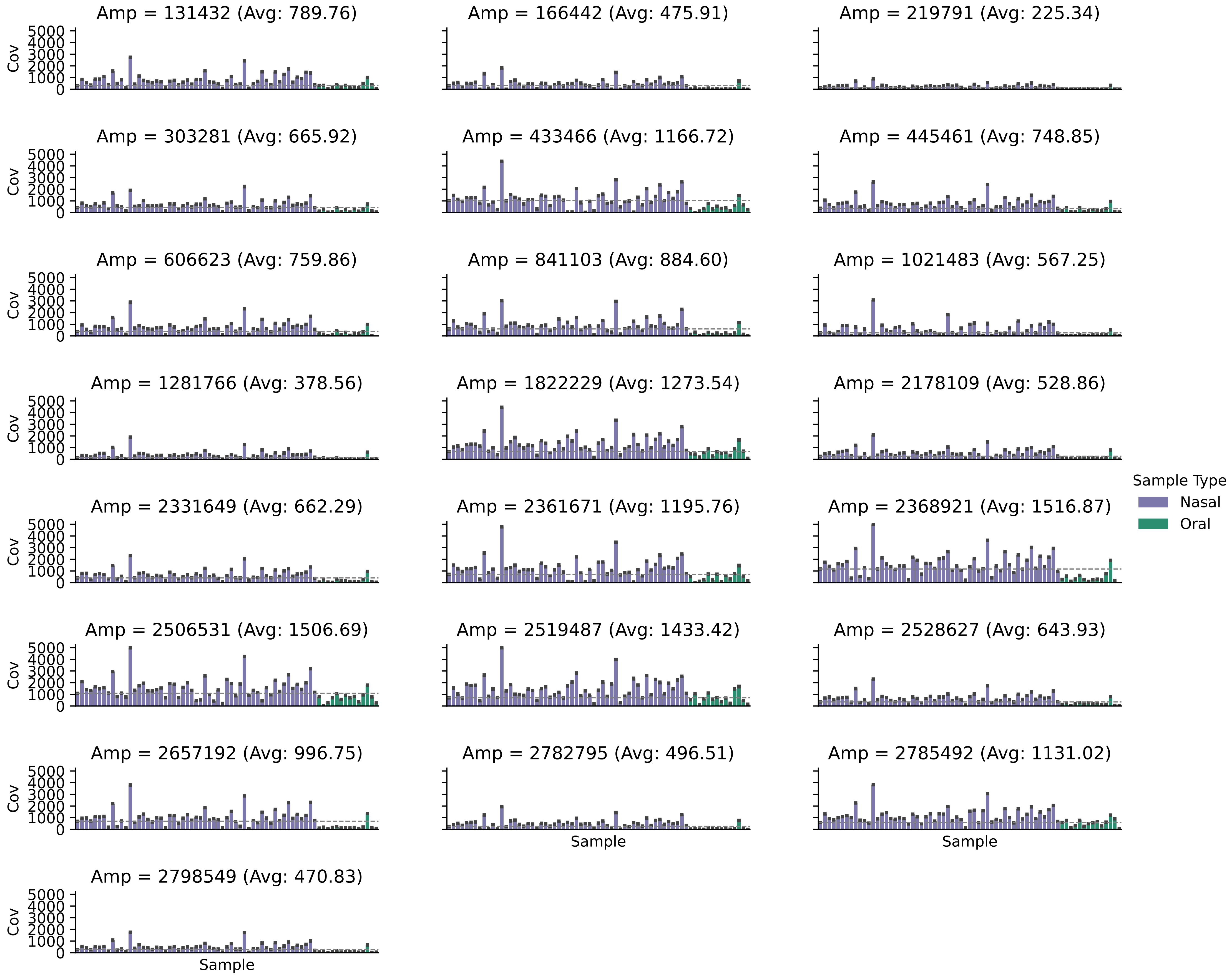
